# Impact of Enteric Neuronal Loss on Intestinal Cell Composition

**DOI:** 10.1101/2024.06.26.600730

**Authors:** Naomi J.M. Kakiailatu, W. Zhang, Laura E. Kuil, Jonathan D. Windster, Eric Bindels, Joke T.M. Zink, Michael Vermeulen, Bianca M. de Graaf, Deepavali Sahadew, Thierry P.P. van den Bosch, Demi Huijgen, Cornelius E.J. Sloots, Rene M.H. Wijnen, Robert M.W. Hofstra, E. de Pater, Veerle Melotte, Maria M. Alves

## Abstract

Hirschsprung disease (HSCR) is a congenital disorder characterized by the absence of an enteric nervous system (ENS) in the distal gut. While the ENS is critical for normal gut function, its broader role in maintaining intestinal homeostasis remains underexplored.

Using single-cell RNA sequencing, we investigated the impact of ENS loss on gut composition in wildtype and *ret* mutant (HSCR model) zebrafish. Significant alterations were identified, including increases in immune cells and shifts in epithelial and extracellular matrix (ECM)-producing cell populations. Immune dysregulation was highlighted by impaired TNF-α signaling via NF-κB, while epithelial cell changes pointed to disrupted energy homeostasis with downregulated fatty acid metabolism and cell cycle pathways. Furthermore, the ECM producing cells showed enriched fibrotic markers. Alterations of the intestinal composition were validated in human HSCR tissues, underscoring the clinical relevance of these findings. These changes can underlie the development of secondary complications and be potentially used to improve patient outcomes.

## Introduction

The enteric nervous system (ENS) is a large network of neurons and glia that not only controls intestinal motility, but is also responsible for a diverse range of functions within the intestine. These include facilitating communication with various intestinal cell types, thereby influencing mucosal transport and secretions. Additionally, the ENS plays a significant role in modulating immune and endocrine responses, as well as regulating blood flow within the gastrointestinal (GI) tract (1). The ENS arises from enteric neural crest cells (ENCCs) which migrate along the gut, proliferate and differentiate to colonize the whole intestine (1, 2). Defects in any of these processes can lead to GI motility disorders associated with a variety of enteric neuropathies (3). Hirschsprung disease (HSCR) is the most common congenital enteric neuropathy, occurring approximately in one in 5.000 newborns (4, 5). HSCR is characterized by the lack of enteric neurons in the colon, most commonly in the distal part, resulting in intestinal constipation and obstruction (1, 6). HSCR is a complex multi-genetic disease and to date, 30 HSCR genes have been identified, which together explain approximately 30% of all cases. However, the most important gene involved in HSCR pathogenesis is the Rearranged during transfection gene (*RET*). *RET* codes for a transmembrane tyrosine kinase receptor that is expressed in ENCCs, supporting their survival, proliferation, differentiation, and migration (7, 8). Pathogenic variants in *RET* account for 50% of familial and 15-20% of sporadic cases (9, 10).

Standard treatment for HSCR is pull-through surgery, in which the aganglionic part of the colon is removed. This surgery is performed in the first few months of life (11-13). Although this procedure is life-saving, it is not curative and is accompanied by secondary complications in 32% of the patients (14). Early complications include anastomotic leak, enterocolitis, perianal excoriation and adhesional obstruction. Long-term complications are associated with chronic obstructive symptoms, enterocolitis and soiling. Furthermore, up to 10% of children diagnosed with HSCR may require a colostomy or further surgery to treat constipation or incontinence. These clinical symptoms strongly indicate a persistent problem in the ganglionic region of the colon and rectum, which remains even after surgical removal of the affected bowel segment (15, 16). Such observations not only suggest a possible impairment of the ENS in the ganglionic bowel, but also a potential effect on other intestinal cell types. Despite significant advancements, most existing studies have only focused on the ENS. For instance, single-cell RNA sequencing (scRNA-seq) on *ret*-deficient mice has provided valuable insights into cellular interactions and pathways disrupted in disease (17). However, the broader impact of ENS absence on other intestinal cell types is often overlooked, leaving gaps in our understanding of how ENS deficiency affects intestinal composition during HSCR pathogenesis.

Zebrafish provide unique advantages for studying HSCR due to their genetic similarity to humans, the optical transparency of their embryos, and the conservation of key developmental pathways involved in ENS formation (18-21). Additionally, their suitability for high-throughput genetic screening enables the identification of candidate genes involved in HSCR by analyzing their roles in ENS development (20). The *ret* mutant zebrafish is particularly valuable for ENS research, as it shows a complete absence of enteric neurons in the intestine, closely mirroring the colonic phenotype observed in HSCR patients (19, 22). Furthermore, the availability of transgenic lines, such as the tg(*phox2bb*:GFP) line, which fluorescently labels enteric neurons, enables detailed visualization and analysis (23). Together, these features make the zebrafish an outstanding model for studying the total intestinal composition in the absence of an ENS. In this study, we use the zebrafish to investigate which intestinal cell types are affected due to absence of the ENS. For this, we performed single cell RNA sequencing (scRNA-seq) on isolated intestines from both wildtype and *ret* mutant zebrafish, with a total colonic HSCR phenotype. Several cellular changes were identified, which might account for the secondary complications diagnosed in HSCR patients, and may even contribute to future treatment options.

## Results

### Differential cellular composition in *ret* mutant zebrafish intestine

To explore the impact of enteric neuronal loss on other, non-neuronal, intestinal cell types, we performed scRNA-seq on 5 days post fertilization (dpf) zebrafish intestines. We analyzed both wildtype tg(*phox2bb*:GFP) and *ret^hu2846/hu2846^* tg(*phox2bb*:GFP) zebrafish, referred here as *ret* mutant, which were selected for their total colonic HSCR phenotype (Figure 1A) (22). For the wildtype group, 244 intestines were isolated, yielding 9.010 cells after filtering out low quality droplets, while for the *ret mutant,* 266 intestines were collected, yielding 10.347 cells. Datasets obtained were further integrated by the standard Seuratv3 workflow and normalized by SCTransform, leading to the identification of 8 distinct cell clusters, namely, epithelial cells (*epcam*), endothelial cells (*kdrl, flt1*), ENS cells (*phox2bb, elavl3, sox10*), immune cells (*coro1a, lcp1, ccl36.1*), muscle cells (*acta2, mylkb*), erythroid cells (*hbbe1.1, hbbe1.3, hbae3)*, thrombocytes (*itga2b, gp1bb*), as well as cells enriched in extracellular matrix (ECM) components such as *col1a2*, *col1a1*, and *cxcl12a* (Figure 1B,C and Data S1). Interestingly, *ret* expression was still observed in the *ret* mutant dataset, specifically in the ENS and epithelial clusters, suggesting that a subset of the total colonic *ret* mutant fish sequenced might actually be *ret* heterozygous (*ret*^+/-^) zebrafish (Figure 1D). This result was not surprising, considering the high phenotypic variability described in *ret*+/- zebrafish (22).

**Figure 1.**
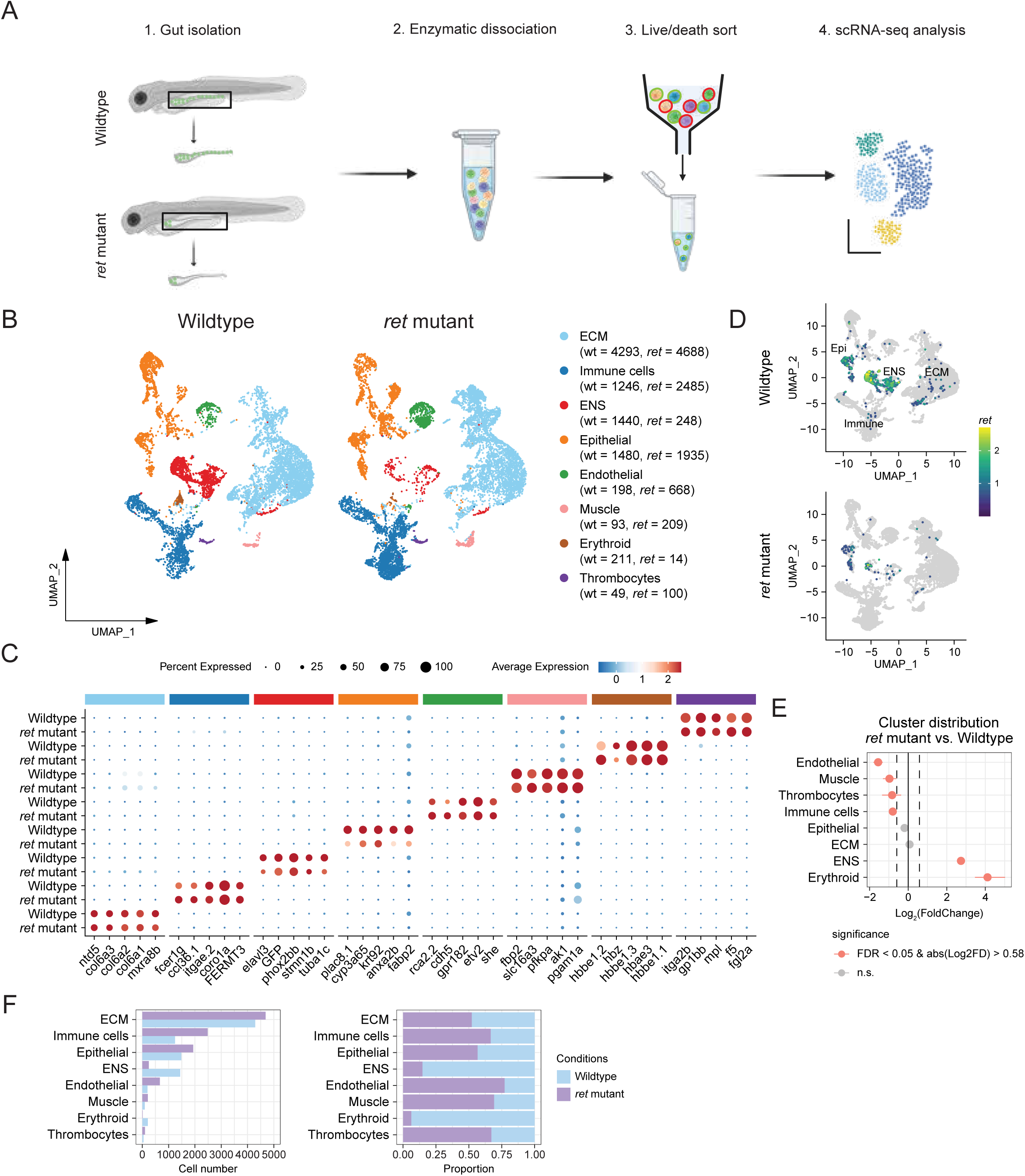
Single-Cell transcriptomic profile of zebrafish intestinal cells in wildtype and *ret* mutants. (A) Experimental design of isolated zebrafish intestines for scRNA-seq. Intestines of 5 dpf wildtype tg(*phox2bb*:GFP) and *ret* mutant tg(*phox2bb*:GFP) zebrafish were isolated and enzymatically dissociated. FACS was used to sort live, single cells for single-cell RNA-sequencing (scRNA-seq). (B) Uniform manifold approximation and projection (UMAP) analysis of scRNA-seq showing 8 distinct cell types. UMAP atlas was separated between control (wildtype) and *ret* mutant zebrafish cells. The colored bar on the top indicates subclusters as shown in (A) and features were separated according to subclusters. (C) Feature dot plot representing the expression level of top 5 transcripts (ordered by average log_2_ fold change) per cluster between wildtype and *ret* mutant zebrafish cells. Features were separated according to clusters. (D) Expression pattern of *ret* in UMAP atlas, separating between wildtype and *ret* mutant zebrafish samples. Navy-blue indicates lower expression levels, while yellow indicates higher expression levels. Grey shows non *ret* expressing cells. (E) Cluster size comparison of each cell type between wildtype and *ret* mutant zebrafish cells. Significantly differentially distributed clusters are represented in orange. The significance of differences was determined by threshold of the false discovery rate (FDR) < 0.05 and absolute log_2_ fold change > 0.58. The dotted line indicates the threshold of log_2_ fold change. (F) Bar plot (left panel) and stacked bar plot (right panel) showing the cell number contributing to each cluster for all cell types in control (wildtype, light-blue) and *ret* mutant zebrafish (light-purple) samples.

Proportion analysis revealed significant shifts in cell populations between wildtype and *ret* mutant zebrafish intestines (Figure 1E, F). As expected, a significant reduction in ENS cells was observed in the *ret* mutant zebrafish (Figure 1E, F). However, our analysis also indicated that other intestinal cell types were affected. To further explore the specific effects on different cell types, we performed subset analysis focusing on the ENS, immune, epithelial and ECM clusters. While an increase in endothelial and muscle cells, and a decrease in erythroid cells was also detected, no further sub-analysis was performed on these groups due to the relatively low number of cells captured.

### ENS loss in *ret* mutant zebrafish

Sub-analysis of the ENS cluster identified eleven distinct subclusters (Figure 2A, B). These included Schwann cell progenitors (SCPs) (*mmp17b, clic6, sox10*), neural crest progenitors (*ncam1b, hoxb5b*), proliferating cells (*cdk1*, *mki67*, *pcna*), migrating neural crest cells (*dkk1b*, *emp3b*), differentiating enteric neurons (*phox2bb, phox2a, myt1b*), and *notch* responsive differentiating neurons (*her4.2.1, notch1a, notch3*) (24). Additionally, we identified enteric glia (*cx43, sox2, her4.1*), and four types of differentiated neurons including, inhibitory motor neurons (*vipb, nos1)*, motor neurons (*olig2*, *isl2b*), sensory intrinsic primary afferent neurons (IPANs) *(nmu, vgf, tac3a and calb2a*) and *phox2bb* negative neurons (*elavl3, gad1b, sv2a*) (24) (Figure 2B and Data S2). *ret* expression could be detected in all subclusters, except in SCPs, differentiating neurons, IPANs and enteric glia (Figure 2C). Analysis of the ENS cluster in the *ret mutant* zebrafish confirmed previous results, showing that nearly all neural subclusters were affected, except for SCPs (Figure 2D, E). A more detailed characterization of the ENS changes observed in the *ret* mutant zebrafish has been previously described by us elsewhere (25).

**Figure 2.**
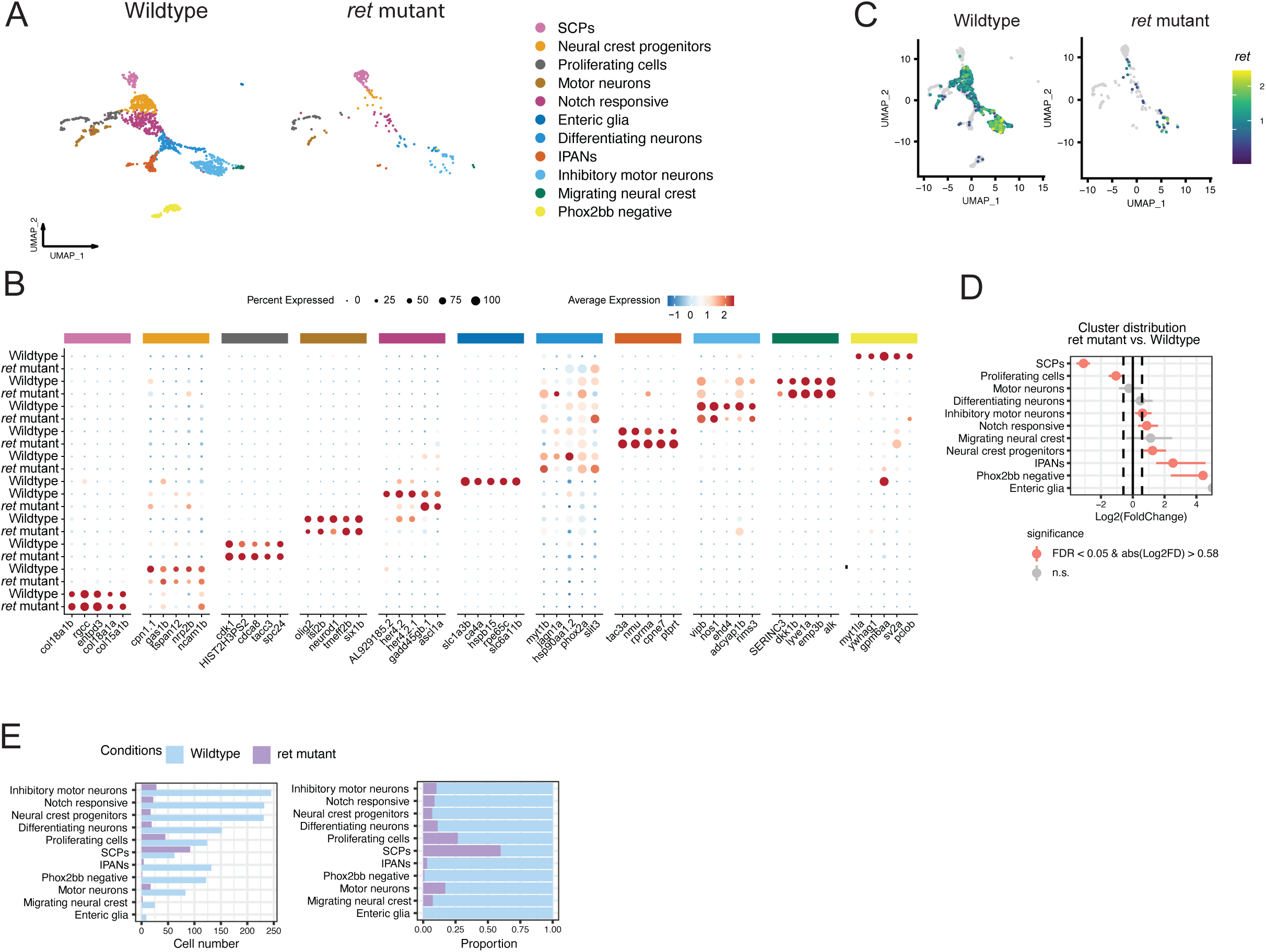
Molecular features of ENS in *ret* mutant zebrafish. (A) Uniform manifold approximation and projection (UMAP) analysis of enteric nervous system (ENS) cells showing all 11 subclusters. UMAP atlas was separated between wildtype and *ret* mutant zebrafish cells. (B) Feature dot plot representing the expression level of top 5 transcripts (ordered by average log_2_ fold change) per subcluster of ENS between wildtype and *ret* mutant zebrafish cells. The colored bar on the top indicates subclusters as showing in (A) and features were separated according to subclusters. (C) The expression pattern of *ret* in UMAP atlas of ENS, separating between wildtype and *ret* mutant zebrafish samples. Navy-blue indicates lower expression levels, while yellow indicates higher expression levels. Grey shows non *ret* expression cells. (D) Cluster size comparison of each ENS subclusters between wildtype and *ret* mutant zebrafish cells. Significantly differentially distributed subclusters are represented in orange. The significance of differences was determined by threshold of the false discovery rate (FDR) < 0.05 and absolute log_2_ fold change > 0.58. Dotted line indicates the threshold of log_2_ fold change. (E) Bar plot (left panel) and stacked bar plot (right panel) showing the subcluster size number of ENS contributing to each subcluster of all ENS in wildtype (light-blue) and *ret* mutant zebrafish (light-purple) samples.

### Increase of immune cell populations in *ret* mutant zebrafish

In zebrafish, innate immune cells begin to emerge as early as 15 hours post-fertilization (hpf), providing the first line of defense. These cells, such as macrophages and neutrophils, are fully functional well before the development of the adaptive immune system. In contrast, adaptive immune cells, including T cells, start to appear at 5 dpf, while B cells appear at 20 dpf (26). In the subset analysis of immune cells, five subclusters were identified (Figure 3A). These included a group of proliferating immune cells (*csf1rb*, *rpl13,* and *dnmt3bb*) and immune progenitors (*coro1a and ccr9a*) (27-31). Other subclusters observed consisted of macrophages (*mpeg* and *mfap4*) *(32)*, T/NK cells (*ccl34b.1*, *mef2b*, *sash1b*, and *runx3*a), and a small cluster of innate lymphoid cells type 2 ( *il4* and *gata3*) (31, 33) (Figure 3B and Data S3). Only a neglectable number of cells showed *ret* expression (Figure 1D). Comparison of the immune clusters of wildtype and *ret* mutant zebrafish, showed an increase in cell proportions of all immune cells in the *ret* mutant (1246 vs 2485), except of macrophages (Figure 3C, D).

**Figure 3.**
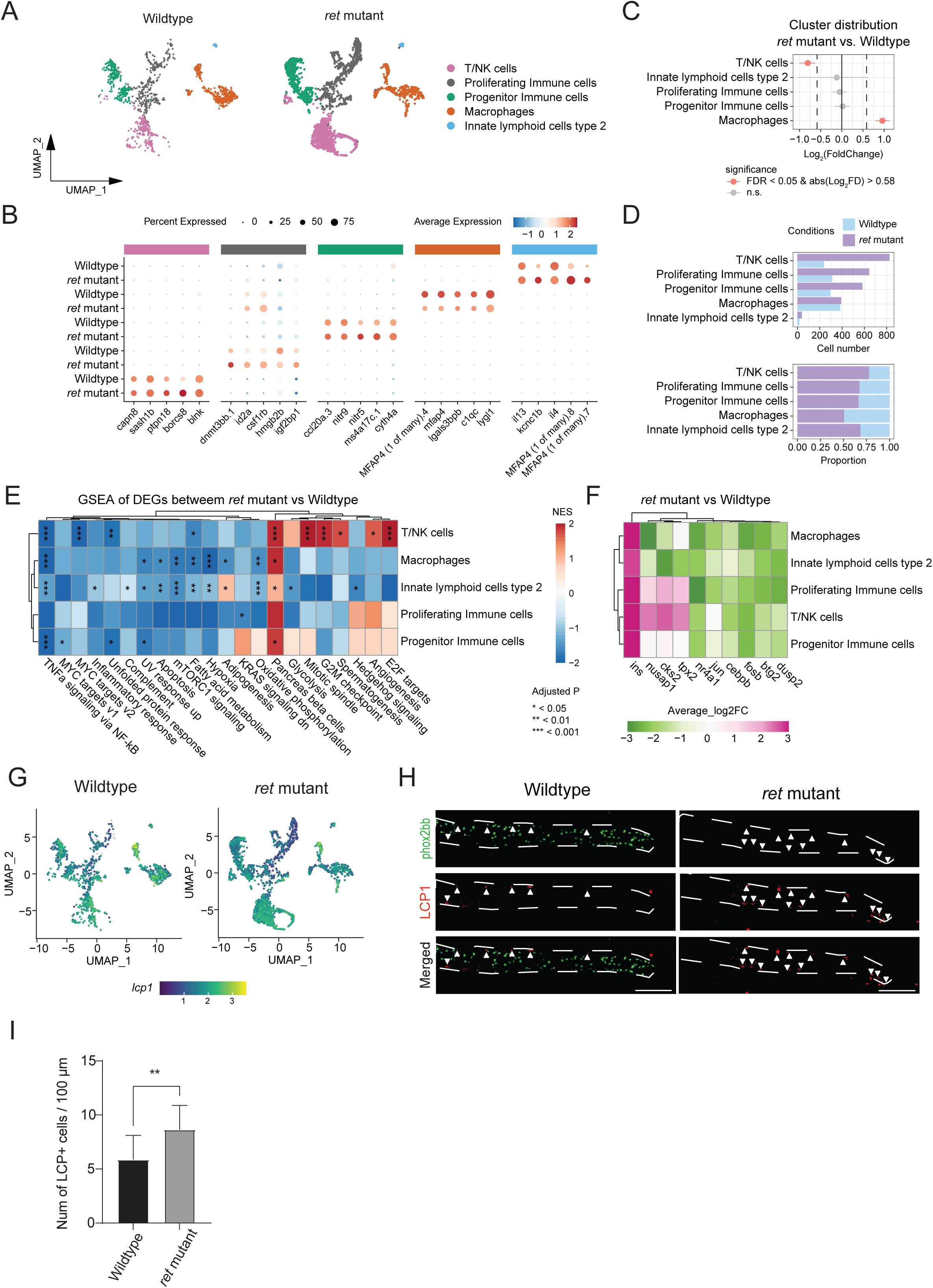
Unique functionality of immune cells in *ret* mutant zebrafish. (A) Uniform manifold approximation and projection (UMAP) analysis of immune cells showing all 5 subclusters. UMAP atlas was separated between wildtype and *ret* mutant zebrafish cells. (B) Feature dot plot representing the expression level of top 5 expressed (ordered by average log_2_ fold change) transcripts per subcluster of immune cells between wildtype) and *ret* mutant zebrafish cells. The colored bar on the top indicates the subclusters as shown in (A) and features were separated according to subclusters. (C) Cluster size comparison of each immune cell subcluster between wildtype and *ret* mutant zebrafish cells. Significantly differentially distributed subclusters represented in orange. The significance of differences was determined by threshold of the false discovery rate (FDR) < 0.05 and absolute log_2_ fold change > 0.58. The dotted line indicated the threshold of log_2_ fold change. (D) Bar plot (top panel) and stacked bar plot (bottom panel) showing the cell number contributing to each subcluster of all immune cells in wildtype(light-blue) and *ret* mutant zebrafish (light-purple) samples. (E) Gene set enrichment analysis (GSEA) of hallmark gene signatures representing different functionality of immune cells subclusters (77). Blue indicates the down-regulated pathways in *ret* mutant zebrafish immune cells subclusters and red indicates the up-regulated pathways in *ret* mutant zebrafish immune cells. The significance threshold of FDR adjusted P value (Adjusted P) < 0.05 was applied. NES: Normalized enrichment score. (F) Heatmap of differentially expressed genes, which were involved in significantly altered hallmark pathways between wildtype and *ret* mutant immune cells subclusters. Green indicates the down-regulated gene in *ret* mutant immune cells subclusters while pink indicates the up-regulated gene in *ret* mutant immune cells subclusters. (G) UMAP atlas showing the expression pattern of *lcp1* in immune cells, separating between wildtype) and *ret* mutant zebrafish samples. Navy-blue indicates lower expression levels, while yellow indicates higher expression level. Grey shows non *ret* expression cells. (H) IHC staining of LCP1 in tg(*phoxbb*:GFP) wildtype and *ret* mutant zebrafish, showing increased numbers of LCP1+ cells in the *ret* mutant zebrafish. Representative maximum projections from 5 dpf larvae. The intestine is marked by the white dashed lines. Phox2bb+ cells are shown in green and LCP1+ cells in red, depicted by the white arrowheads. Scale bar represents 80 μm. (I) Graph showing significant increased numbers of LCP1+ cells in *ret* mutant zebrafish (p=0,005) (n=12 per group).

Gene Set Enrichment Analysis (GSEA) revealed that TNF-α signaling via NF-κB, was significantly downregulated in the *ret* mutant across almost all immune cell clusters, except for the innate lymphoid cells type 2, pointing to a potential dysfunction of TNF-α signaling in the absence of *ret* expression (Figure 3E). This downregulation was further supported by reduced expression of specific genes involved in immune response and inflammation, including *nr4a1, cebpb, fosb* and *dusp2* (Figure 3F). Interestingly, we also observed a significant upregulation of cell cycle-related pathways, particularly in T/NK cells, indicating heightened proliferative activity of all immune populations (Figure 3E). Notably, *nusap1*, *cks2*, and *tpx2* were specifically upregulated in proliferating immune cells and T/NK cells (Figure 3F).

To validate the increased number of immune cells in the *ret* mutant, we performed whole-mount immunohistochemistry (IHC) on both wildtype and *ret mutant* zebrafish using L-plastin, a leukocyte marker expressed in all immune clusters (Figure 3G). An increased number of Lcp+ cells was detected in *ret* mutant zebrafish (*p* = 0.005), confirming the scRNA-seq results (Figure 3H, I).

### Changes in epithelial cell types in ret mutant zebrafish

To investigate the impact of the ENS on epithelial cells, a sub-analysis was performed. Nine distinct sub-clusters were identified, including differentiating epithelial cells (*epcam, her15, cotl1*), differentiating enterocytes (*krt4, krt8, sparc, pdx1*), BEST4/OTOP2+ enterocytes (*best4, otop2, cftr*), anterior enterocytes (*fabp2, ada, fabp1b.1, vil1*), and mid-region enterocytes (*fabp6, slc10a2, tmigd1*) (34, 35). A small cluster of enterochromaffin cells (ECCs) expressing *tph1a* and *tph1b* was also observed. In addition, a minor population of goblet cells expressing *agr2* and *sstr5* was identified (36). Notably, *sstr5* was also expressed in the BEST4/OTOP2+ enterocyte population. Furthermore, two distinct clusters of enteroendocrine cells (EECs) were identified: EEC 1 (*neurod1*, *pax6b* and *ccka*) and EEC 2 (*neurod1*, *isl1*, *cpe*, and *pcsk2*)(37-43) (Figure 4A, B and Data S4).

**Figure 4.**
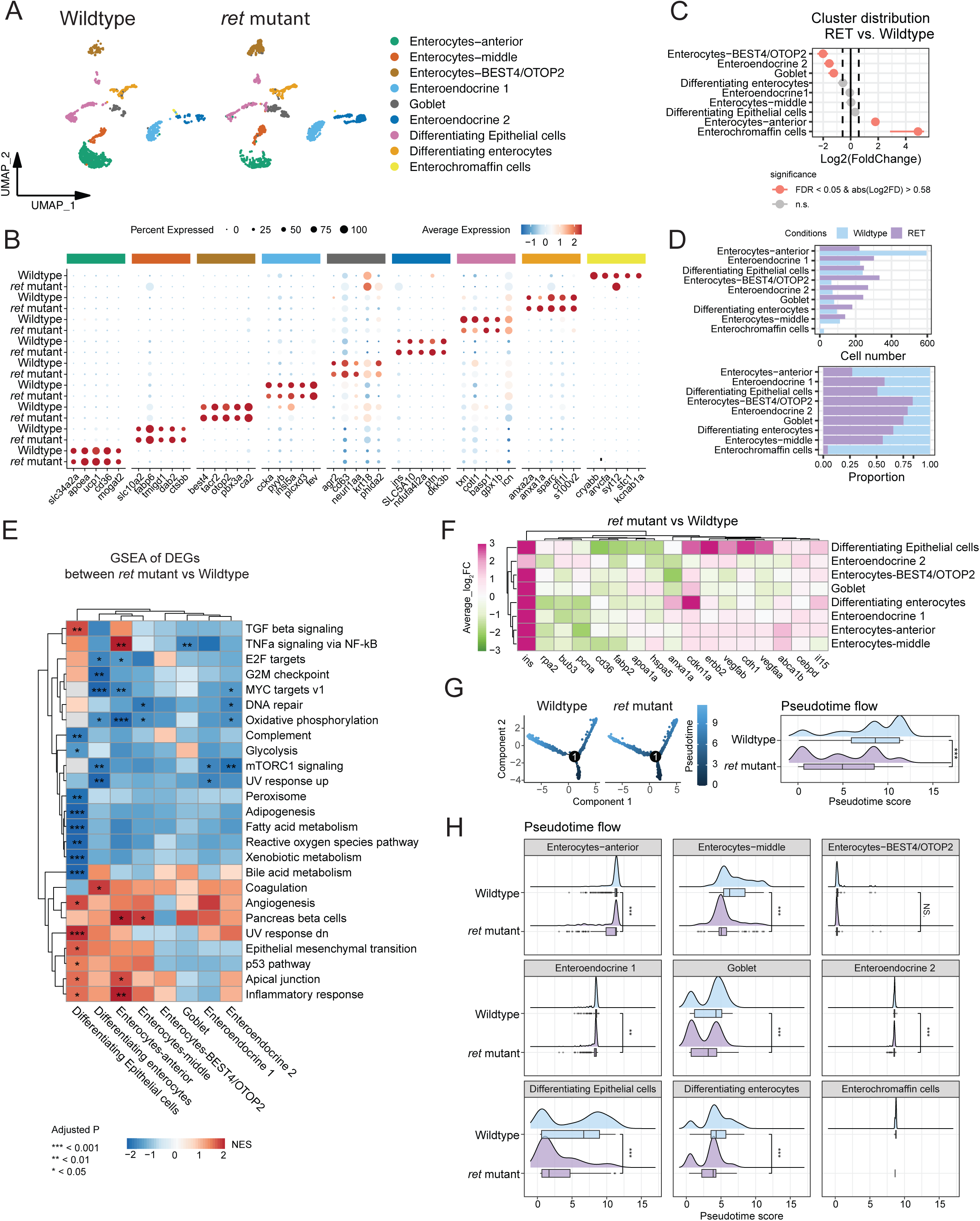
Epithelial subcluster identification and functional pathway shifts in *ret* mutant zebrafish. (A) Uniform manifold approximation and projection (UMAP) analysis of epithelium showing all 9 subclusters. UMAP atlas was separated between wildtype and *ret* mutant zebrafish cells. (B) Feature dot plot representing the expression level of top 5 transcripts (ordered by average log_2_ fold change) per subcluster of epithelium between wildtype and *ret* mutant zebrafish cells. The colored bar on the top indicates subclusters as shown in (A) and features were separated according to subclusters. (C) Cluster size comparison of each epithelium subcluster between wildtype and *ret* mutant zebrafish cells. Significantly differentially distributed subclusters are represented in orange. The significance of differences was determined by the threshold of the false discovery rate (FDR) < 0.05 and absolute log_2_ fold change > 0.58. The dotted line indicated the threshold of log_2_ fold change. (D) Bar plot (top panel) and stacked bar plot (bottom panel) showing the cell numbers contributing to each subcluster of all epithelial cells in wildtype(light-blue) and *ret* mutant zebrafish (light-purple) samples. (E) Gene set enrichment analysis (GSEA) of hallmark gene signatures representing different functionality of epithelium subclusters (77). Blue indicates the down-regulated pathways in *ret* mutant zebrafish epithelium subclusters and red indicates the up-regulated pathways in *ret* mutant zebrafish epithelium. The significance threshold of FDR adjusted P value (Adjusted P) < 0.05 was applied. NES: Normalized enrichment score. Due to only 1 cell contained in *ret* mutant enterochromaffin cells cluster, differentially expressed genes could not be detected, resulting in the enterochromaffin cells cluster being excluded from GSEA approach. (F) Heatmap of differentially expressed genes, which were involved in significantly altered hallmark pathways between wildtype and *ret* mutant epithelium subclusters. Green indicates the down-regulated gene in *ret* mutant epithelium subclusters while pink indicates the up-regulated gene in *ret* mutant epithelium subclusters. (G) Pseudotime trajectories (left panel) showing the developmental track of epithelium involved in wildtype) and *ret* mutant zebrafish intestines. Dark-blue indicates lower pseudotime scores, light-blue indicates higher pseudotime scores, and number within trajectories indicates the branch node. Ridge chart (right panel) showing the distribution of epithelium along with pseudotime trajectory (left panel), separating between wildtype and *ret* mutant zebrafish. Box chart showing the statistical pseudotime score analysis between wildtype and ret mutant epithelium. T-test was used to calculate p value. (H) Subcluster-based individual ridge charts showing the distribution of each epithelium subcluster along with pseudotime trajectory as shown in (G) right panel, separating between wildtype and ret mutant samples. Box chart showing the statistical pseudotime score analysis between wildtype and ret mutant epithelium. t-test was used to calculate p value. NS.: Not significant; **: P < 0.01; ***: P < 0.001.

In the *ret* mutant zebrafish, we observed significant alterations in the epithelial cell populations, compared to wildtype zebrafish. The most notable findings included a significant reduction in anterior enterocytes (595 vs 221) and ECCs (29 vs 1), contrasted by an increase in EEC2 (71 vs 269), goblet cells (80 vs 242), and BEST4/OTOP2+ enterocytes (64 vs 333) (Figure 4C, D). GSEA revealed distinct alterations in the *ret* mutant zebrafish, especially in metabolic and signaling pathways within the epithelial cell populations. Notably, fatty acid metabolism was significantly downregulated in both anterior enterocytes and differentiating epithelial cells, specifically marked by overall reductions in *apoa1a* and *fabp2* (Figure 4E, F). This downregulation can have implications for cellular energy homeostasis and overall epithelial function. In contrast, inflammatory response pathways were markedly upregulated, particularly those linked to TNFα signaling via NF-κB, with notable increases in *cdkn1a* and *cebpd* (Figure 4E, F). This indicates an elevated inflammatory state within the epithelium. Additionally, a marked decrease in cell cycle pathway activity was observed in the differentiating enterocytes cluster, as indicated by overall lower expression of *pcna* and *rpa2* (Figure 4E, F). Such downregulation likely reflects disruptions in normal cell proliferation and differentiation, potentially compromising tissue regeneration and maintenance. To further understand how the loss of ENS impacts epithelial differentiation and maturation, a pseudotime analysis was performed, revealing three distinct branches corresponding to enteroendocrine cells, enterocytes, and BEST4+ enterocytes/goblet cells (Supplementary Figure 1). Interestingly, the overall pseudotime trajectory is significantly reduced in the *ret* mutant (Figure 4G). Pseudotime progression within each epithelial cluster, was also decreased across all clusters, except the BEST4+ enterocyte cluster, with the most pronounced reductions occurring in the differentiating epithelial and differentiating enterocyte clusters (Figure 4H). These findings suggest that the loss of ENS specifically interferes with the maturation of epithelial progenitors into functional cell types, which could have downstream effects on intestinal homeostasis and function.

Profound alterations in ECCs, BEST4/OTOP2+ enterocytes, and anterior enterocytes, were validated by IHC and fluorescent *in situ* hybridization (FISH). Reduction in ECCs was confirmed by IHC using a serotonin (5-HT) antibody. Considering that 5-HT-is also expressed in ENS cells, only *phox2bb*-negative cells were considered to be ECCs. Using this approach, we observed a significant loss of ECCs in the ret mutant intestines, that was independent of the loss of ENS cells (p=3.518E-05) (Figure 5A, B). However, future co-staining with a more specific ECC marker could enhance precision of this finding. FISH analysis for *best4* also corroborated the scRNA-seq results, demonstrating a marked increase in *best4*+ enterocytes in the ret mutant zebrafish (p=0,025) (Figure 5C, D). This increase was most striking in the distal intestine, where *best4*+ enterocytes were entirely absent in wildtype zebrafish but prominently present in the *ret* mutant (p=0,021). Finally, reduction in anterior enterocytes was confirmed through *in situ* hybridization using a *fabp2* probe, which revealed a decreased expression in the *ret* mutant intestine (Figure 5E).

**Figure 5.**
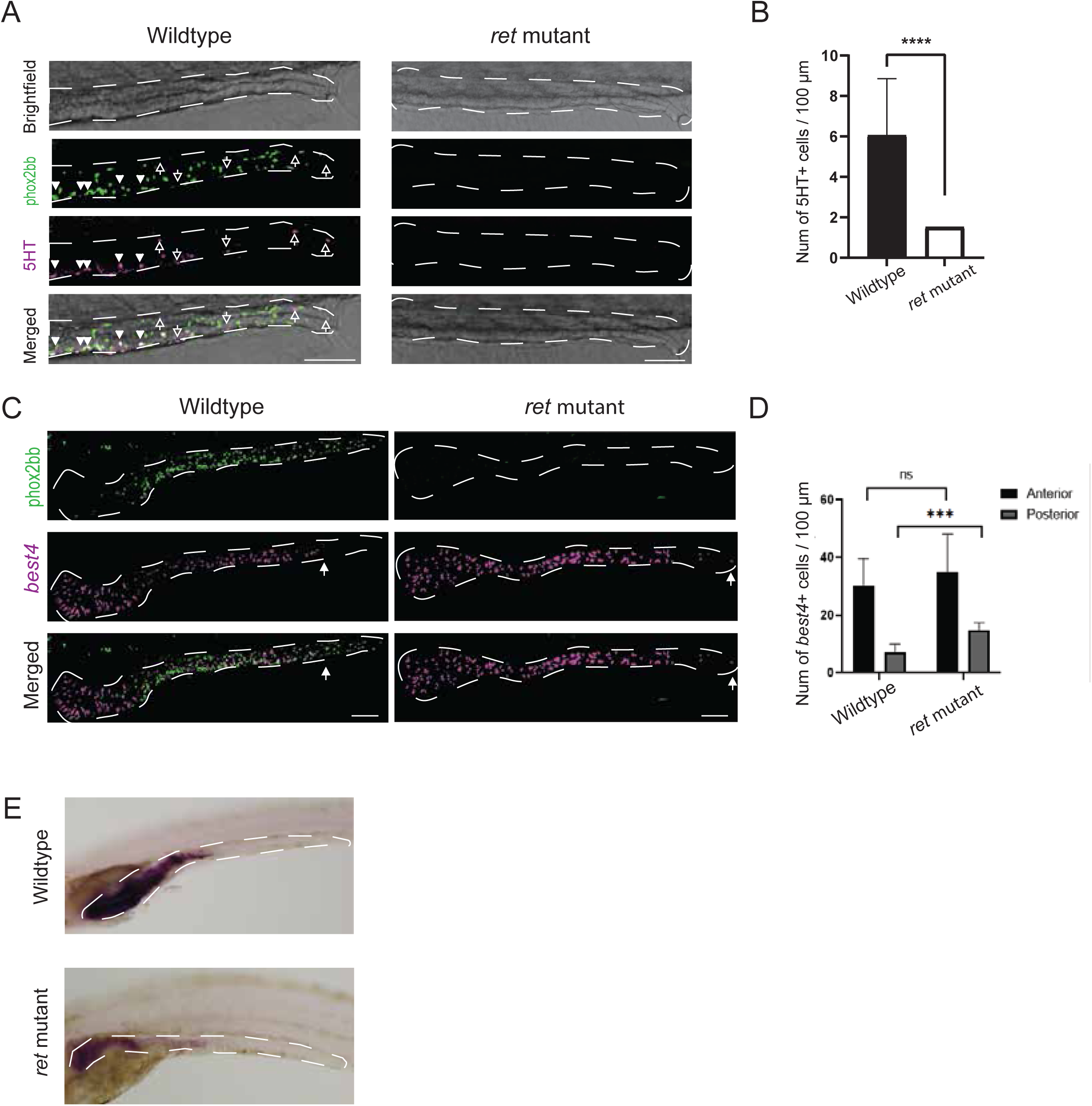
Altered epithelial composition in *ret* mutant zebrafish larvae. A) Maximum projections of immunohistochemistry staining of 5-HT in tg(*phox2bb*:GFP) wildtype (n=14) and *ret* mutant (n=12) zebrafish. Intestines are marked by the white dashed lines. Phox2bb+ cells are shown in green and 5-HT+ cells in magenta. Scale bar represents 80 μm. B) Graph showing decreased numbers of 5-HT+ cells in ret mutant zebrafish (p=3,09918E-05). C) Fluorescent *in situ* hybridization of *best4* in tg(*phox2bb*:GFP) wildtype and *ret* mutant zebrafish. Representative maximum projections from 5 dpf wildtype and ret mutant tg(*phox2bb*:GFP) zebrafish. Intestines are marked by the white dashed lines. Phox2bb+ cells are highlighted in green and *best4*+ cells in magenta. Arrows indicate the presence of Best4+ enterocytes. D) Quantification of the number of *best4*+ cells in both anterior (p=0,128) and posterior regions (p=0,021) of the intestine, in wildtype (n=10) and ret mutant zebrafish (n=6). E) In situ hybridization of *fabp2* in wildtype (n=10) and *ret* mutant (n=10) zebrafish, scale bar represents 100 μm.

### ECM Sub-Analysis Reveals Fibrosis-Associated Signaling in the ret mutant zebrafish

Sub-analysis of the ECM expressing cells revealed 8 distinct subclusters (Figure 6A), consisting of immature ECM-producing cells (*vim, hoxb5a, rpl7a, rpl12* genes), differentiating myofibroblasts (*pnf1, tubb2b, rpl genes*), myofibroblasts (*myl9, fn1*) and three distinct fibroblast subclusters. Fibroblasts 1 expressed collagen genes (*col6a2*, *col6a4a*) and transcription factors (*foxf2a*, *twist1b)*, indicating their crucial role in maintaining structural integrity and supporting tissue repair and regeneration (44). Fibroblasts 2 feature genes like *cav1* and *jam2b*, which are associated with cellular signaling and maintaining cell junctions integrity (45). These fibroblasts are essential for the function of the intestinal barrier and its adaptive response to environmental stimuli. Fibroblasts 3 are marked by a diverse array of collagen genes and remodeling enzymes, such as *mmp2*, highlighting their involvement in ECM remodeling needed for tissue growth and healing, in response to physical and inflammatory stress (46). Additionally, two clusters of mesenchymal cells were identified, one consisting of differentiating mesenchymal cells (*hox* genes, *col12a1a, thbs4b*), and a smaller cluster of mature mesenchymal cells (*itga10, bgna, ssp1, bmp8a*) (Figure 6A, B and Data S5). Comparing the ECM clusters in the *ret* mutant with the wildtype, showed a significant increase in fibroblasts 2 (185 vs 1072) and mesenchymal cells (5 vs 130), while a significant decrease was observed in fibroblasts 3 (249 vs 74) and differentiating mesenchymal cells (36 vs 71) (Figure 6C, D). GSEA revealed significant alterations in the ret mutant zebrafish, particularly in metabolic and signaling pathways (Figure 6E). Glycolysis and hypoxia pathways were notably upregulated in differentiating mesenchymal cells, which was further supported by increased expression of glycolysis-related genes (*aldob, eno1a*) and hypoxia-responsive genes (h*mox1a, tgfbi*) (Figure 6F). Additionally, ECM remodeling was evident through the upregulation of key ECM-related genes (*fn1b*, *col18a1a*, and *podxl*), highlighting enhanced structural reorganization and cellular interactions within the ECM (Figure 6F). In contrast, pathways associated with cell proliferation, such as E2F targets and G2M checkpoint, were significantly downregulated across most clusters, with the exception of differentiating myofibroblasts, myofibroblasts, and fibroblasts 3 (Figure 6E). This was reflected in the reduced expression of proliferation-related genes (p*cna, mcm4, mcm5, stmn1a*), indicating a shift away from cell cycle activity towards ECM-focused remodeling and metabolic adaptation (Figure 6F).

**Figure 6.**
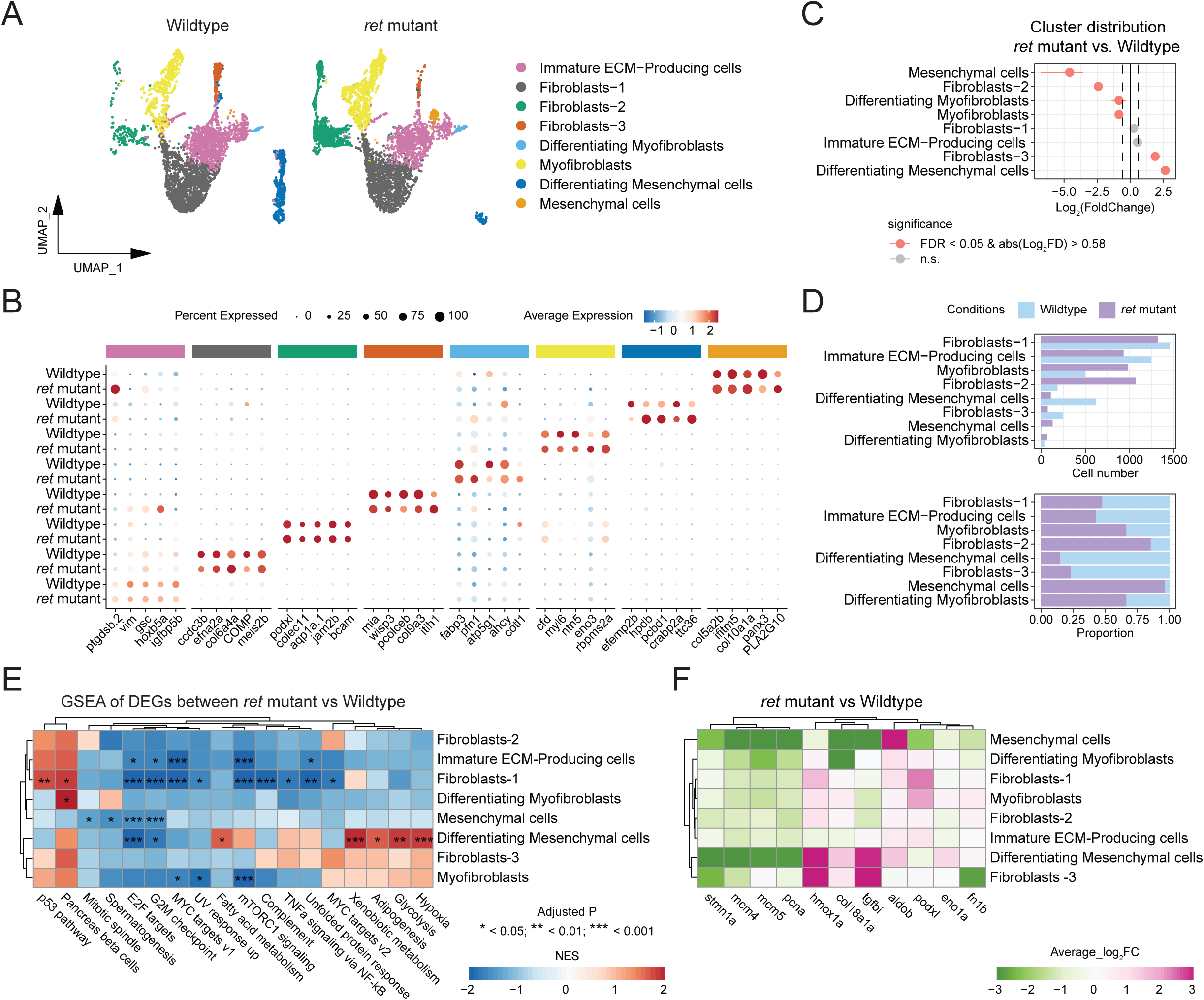
ECM subcluster profiling and intercellular communication dynamics in wildtype and *ret* mutant zebrafish. (A) Uniform manifold approximation and projection (UMAP) analysis of extracellular matrix (ECM) showing all 8 subclusters. UMAP atlas was separated between wildtype and *ret* mutant zebrafish cells. (B) Feature dot plot representing the expression level of top 5 transcripts (ordered by average log_2_ fold change) per subcluster of ECM between wildtype and *ret* mutant zebrafish cells. The colored bar on the top indicates subclusters as shown in (A) and features were separated according to subclusters. (C) Cluster size comparison of each ECM subcluster between wildtype and *ret* mutant zebrafish cells. Significantly differentially distributed subclusters represented in orange. The significance of differences was determined by threshold of the false discovery rate (FDR) < 0.05 and absolute log_2_ fold change > 0.58. The dotted line indicated the threshold of log_2_ fold change. (D) Bar plot (top panel) and stacked bar plot (bottom panel) showing the cell number contributing to each subcluster of all ECM cells in wildtype (light-blue) and *ret* mutant zebrafish (light-purple) samples. (E) GSEA of hallmark gene signatures representing different functionality of epithelium subclusters (77). Blue indicates the down-regulated pathways in *ret* mutant zebrafish ECM subclusters and red indicates the up-regulated pathways in *ret* mutant zebrafish ECM. The significance threshold of FDR adjusted P value (Adjusted P) < 0.05 was applied. NES: Normalized enrichment score. (F) Heatmap of differentially expressed genes, which were involved in significantly altered hallmark pathways between wildtype and *ret* mutant ECM subclusters. Green indicates the down-regulated genes in *ret* mutant ECM subclusters while pink indicates the up-regulated genes in *ret* mutant ECM subclusters.

#### Translational Relevance: Human validations

To assess the translational relevance of our findings in zebrafish to humans, we conducted IHC analyses, targeting the specific cell types affected in our *ret* mutant model: ECCs, immune cells and BEST4+ enterocytes. Human intestinal biopsies collected from a cohort of five controls and five HSCR patients, were used. All patients carried a *RET* mutation and were clinically diagnosed with total colonic aganglionosis, resembling the *ret* zebrafish model used in this study. Control patients had no ENS abnormalities and tissue was taken from visually normal and non-inflamed regions (Table S1).

Immunostainings for 5-HT, a marker of serotonin produced by ECCs, showed a statistically significant decrease of ECCs in HSCR patients, when compared to healthy controls (*p*= 0.0027), (Figure 7A, B). Additionally, immunofluorescence analysis targeting CD45, a pan-leukocyte marker specific of cells in both the innate and adaptive immune systems, demonstrated a substantial increase in CD45+ immune cells within the HSCR patient cohort (*p* = 0.008), (Figure 7C, D). These results align with our zebrafish model outcomes. We also assessed the presence of BEST4+ enterocytes through quantification via IHC, and our results revealed a decrease in the number of BEST4+ cells in the aganglionic bowel of HSCR patients, compared to controls (Figure 7E, F). This result was opposite to the one observed with the *ret* mutant model, as an increase of Best4/Otop2 enterocytes was observed.

**Figure 7.**
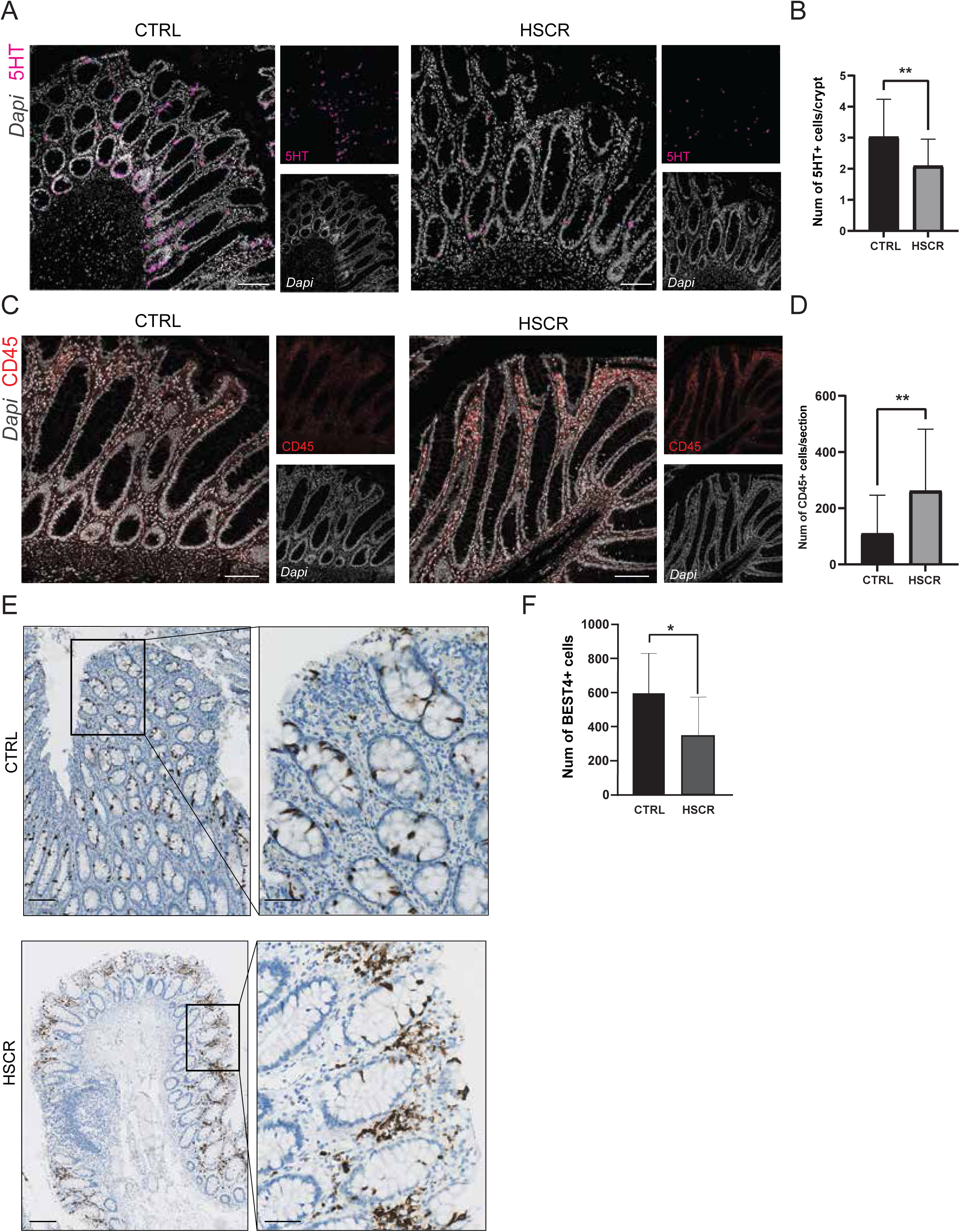
Immunohistochemical validation and quantification in tissue collected from HSCR patients. (A) IHC validation of 5-HT in control and HSCR aganglionic tissue. DAPI is shown in grey and 5-HT+ cells in magenta. Scale bar represents 100 μm. (B) Graph showing significant decrease of 5-HT+ cells per crypt in aganglionic tissue of HSCR patients. (C) IHC validation of CD45+ cells in control and HSCR aganglionic tissue. DAPI is depicted in grey and CD45+ cells in red. Scale bar represents 100 μm. (D) Graph showing significant increase of CD45+ cells in HSCR aganglionic tissue. (E) DAB staining for BEST4 in control and HSCR aganglionic tissue, with zoomed-in views on the right panel. Scale bar represents 200 μm. (F) Graph showing significant decrease of BEST4+ enterocytes in aganglionic tissue of HSCR patients.

## Discussion

Here, we present the single-cell transcriptomic profile of 5 dpf zebrafish intestines, where we identified a diverse range of intestinal cell types including immune, endothelial, muscle, epithelial, and ENS cells. Importantly, by comparing wildtype intestines with those of *ret* mutant zebrafish presenting with a phenotype reminiscent of total colonic aganglionosis in humans, distinct alterations in intestinal composition were observed, which were validated *in vivo*. We further extended our findings beyond the zebrafish to demonstrate their translational relevance to humans, providing additional evidence of an altered human intestinal composition in the absence of an ENS.

As expected, we observed an overall reduction on the number of ENS cells in the *ret* mutant zebrafish, except in the SCPs, which do not express *ret*. This result was consistent with recent findings in *ret* loss-of-function mice (17). Interestingly, the absence of ENS cells was accompanied by significant changes in other cell populations. A significant increase in immune cells was detected in the intestines of *ret* mutant zebrafish, even at an early developmental stage and in the absence of external stimuli, such as feeding. A similar increase in immune cells was observed in intestinal tissue from HSCR patients, suggesting that ENS absence contributes to an imbalanced immune response, potentially influencing the onset of Hirschsprung associated enterocolitis (HAEC). Additionally, we observed a notable downregulation of TNF-α signaling via NF-κB across all immune cell clusters in the *ret* mutant, a pathway essential for neurodevelopmental processes, including embryonic neurogenesis and neural progenitor migration(47-50). Evidence from previous studies support our findings, as they showed lower expression of TNF-α, TLR2, and TLR4 in patients with HAEC, as well as decreased NF-κB subunit expression in the aganglionic colon of HSCR patients (51, 52). Our findings also underscore the critical role of disrupted TNF-α and NF-κB signaling in HSCR, highlighting the need for further research into these mechanisms.

Altered epithelial composition was also observed in the *ret* mutant zebrafish intestines, particularly affecting enterocytes and ECCs. Despite ECCs being a small cluster, their reduction in both *ret* mutants and HSCR patients is particularly noteworthy, due to their critical role in regulating 5-HT, a key modulator of gut motility and secretion. *Ret* is expressed in a subset of ECCs, likely explaining the decreased number of these cells (53). However, it’s noteworthy that inhibiting *Ret* kinase activity in adult mice did not affect 5-HT levels, suggesting that the impact of *ret* on ECCs is specific to developmental stages and/or may vary across species (54). The enrichment of BEST4+ enterocytes, specifically in the posterior gut in the *ret* mutant, may further indicate compensatory epithelial changes in the absence of an ENS. However, in HSCR patients, BEST4+ enterocytes were reduced in the aganglionic regions. Since the most prominent changes in the zebrafish were observed in the distal gut, it is possible that some location-specific differences may not be captured in human samples. Furthermore, the absence of rectum tissue from control patients limited our ability to assess this region comprehensively. Validation of the decreased anterior enterocytes observed in zebrafish was also impeded in human tissue due to restricted access to small intestine tissue, particularly in HSCR patients. Without region-matched samples, direct comparisons of anterior enterocyte abundance between healthy controls and HSCR patients remains challenging.

Alongside the changes in cell abundance, significant alterations in epithelial metabolic and signaling pathways were detected in the *ret* mutant zebrafish. Fatty acid metabolism was notably downregulated in anterior enterocytes and differentiating enterocytes, suggesting that ENS disruption impairs cellular energy homeostasis, essential for maintaining epithelial integrity. This metabolic deficit, together with upregulated inflammatory pathways, notably TNFα signaling via NF-κB, indicates a heightened inflammatory state in the epithelium, potentially as a response to epithelial barrier disruptions, as seen in other GI disorders, such as IBD (55, 56). This inflammatory state aligns with the observed increase in immune cell populations identified in the *ret* mutant and HSCR patients. However, activation of these immune cells appears to be impaired, as the TNFα signaling is downregulated, indicating a disrupted immune response within the gut. Such dysfunction may involve a negative feedback loop that suppresses TNFα signaling due to the increased number of immune cells. In contrast, the elevated TNFα signaling via NF-κB in epithelial cells may represent a compensatory mechanism to address immune cell dysfunction, or directly respond to epithelial barrier damage. Reduced cell cycle activity in epithelial cells further suggests impaired renewal and repair, likely affecting the gut’s ability to maintain its epithelial layer. Overall, these findings underscore the essential role of the ENS in regulating epithelial cell metabolism, inflammation, and regeneration, which can potentially contribute to the secondary complications observed in HSCR patients, specifically HAEC.

Notable changes in ECM producing cells were also detected in the *ret* mutant zebrafish, including a significant increase in fibroblasts 2 and mesenchymal cells, while a significant decrease was observed in fibroblasts 3 and differentiating mesenchymal cells. These shifts were accompanied by upregulation of glycolysis and hypoxia pathways in the differentiating mesenchymal cells of *ret* mutant zebrafish, indicating metabolic reprogramming and adaptation to hypoxic conditions commonly associated with fibrosis (57, 58). Interestingly, these findings are consistent with a recent scRNA-seq study showing increased FN1 expression in the aganglionic regions of HSCR patients, further highlighting potential shifts in ECM composition specific to this condition (59-61).

Taken together, our in-depth single-cell transcriptomic analysis of 5 dpf wildtype and *ret* mutant zebrafish intestines, unravels crucial insights not only into the cellular composition of 5dpf zebrafish intestines, but also into the cellular alterations associated with ENS loss. Validation of our findings in intestinal tissue samples from aganglionic regions of HSCR patients carrying a *RET* mutation, underscores the clinical relevance of our study, as it provides valuable insights into the mechanisms associated with the onset of secondary complications developed by HSCR patients, specifically HAEC. Although further functional studies are needed to deepen our understanding of these mechanisms, our findings open up new avenues for potential therapeutic interventions to prevent such complications. Considering the observed changes in cellular composition, therapies that target immune modulation, and/or strategies to optimize epithelial cell composition, such as anti-inflammatory medications and probiotics, could be explored. Importantly, our results could also benefit the improvement of transplantation therapies, as they show that an abnormal by optimizing the intestinal environment is present in the aganglionic region of HSCR patients, suggesting the need of targeted interventions to optimize the intestinal environment.

### Limitations of the study

One limitation of this study is the limited access to location-matched intestinal samples, which restricted direct comparisons between healthy controls and HSCR patients. Additionally, the anatomical simplicity of the zebrafish intestine, which is shorter and lacks a clear separation between the small intestine and colon, means that observations in zebrafish may not always directly correlate with human physiology, as seen with the BEST4+ enterocytes population. Expanding collaborations with medical institutions to secure a wider range of human tissue samples, including location-matched intestinal segments, would significantly enhance this comparative analysis.

## Materials and methods

### Resource availability

#### Lead contact

Further information and requests for reagent may be directed to, and will be fulfilled by Naomi J.M. Kakiailatu or Maria M. Alves

#### Materials availability

This study did not generate new, unique reagents.

### Experimental model and study participant details

#### Animal husbandry

The following zebrafish lines were used: transgenic tg(*phox2bb*:GFP) (23), tg(*kdrl*:mcherry) (62), and *ret*^hu2846/+^ (19). Zebrafish were kept on a 14/10h light/dark cycle. Embryos and larvae were kept in an incubator at 28.5°C in HEPES-buffered E3 medium. For imaging experiments, fish were treated from 24 hpf onwards with 0.2 mM 1-phenyl 2-thiourea (PTU), to inhibit pigmentation. Animal experiments were approved by the Animal Experimentation Committee of the Erasmus MC, Rotterdam.

### Method details

#### Isolation of zebrafish intestines and pre-processing of intestinal cell suspension for single cell RNA sequencing

Intestines of 5 dpf wildtype and *ret* mutant tg(*phox2bb*:GFP) zebrafish were isolated and dissociated as described before (63, 64). In total, 244 intestines of the wildtype and 266 intestines of the *ret* mutant zebrafish, were isolated and dissociated using papain as a dissociation enzyme.

#### Single-Cell RNA Sequencing Data Analysis

##### Quality Control, Data Integration, Dimensionality Reduction, Normalization, and Clustering

Single-cell RNA-sequencing (scRNA-seq) data were processed using the R package Seurat (version 5.1.0) (65). First, quality control metrics were applied to filter out low-quality cells, following standard thresholds for gene counts (> 100 & < 4500), and mitochondrial gene percentage (< 5%), to retain high-confidence single-cell data. For data integration across samples, the standard Seurat v3 integration workflow (66) was employed to minimize batch effects, enabling robust cross-sample comparisons. The integrated dataset underwent dimensionality reduction using Principal Component Analysis (PCA), followed by Uniform Manifold Approximation and Projection (UMAP) for visualization. SCTransform normalization was applied to adjust for sequencing depth and technical variability, facilitating reliable downstream analysis. Cells were clustered using the default Seurat clustering algorithm.

##### Cluster Composition Analysis

Cluster distribution differences between *ret* mutant and WT fish were analyzed using the R package scProportionTest (version 0.0.0.9000) (67), a tool designed to evaluate proportional changes in cluster composition across conditions, providing insights into condition-specific cellular heterogeneity.

##### Cluster Marker Identification

Cluster-specific markers were identified with Seurat’s FindAllMarkers function, which uses a Wilcoxon rank-sum test to find differentially expressed genes between one cluster versus all others. Default settings were applied, except min.pct was set to 0.5 to filter markers present in at least 50% of cells in a cluster, ensuring robust marker identification and reducing noise from low-abundance transcripts.

##### Gene Set Enrichment Analysis (GSEA)

GSEA was performed to identify enriched biological pathways. The pathway reference gene sets were obtained from the msigdbr R package (version 7.5.1), with the Hallmark gene set category for zebrafish selected to focus on essential pathways (68). Enrichment testing was conducted with the fgsea R package (version 1.30.0), using default parameters to calculate enrichment scores for each pathway, thus identifying condition-associated biological processes and pathways (69).

##### Pseudotime trajectory inference

Pseudotime trajectory inference was performed to compare the development progression between wildtype and ret mutant fish. Trajectory construction and pseudotime score calculation were performed using monocle R package (version 2.32.0)(70, 71). The significance of pseudotime score between wildtype and ret mutant was calculated using t test, the significance threshold is P value < 0.05.

#### Zebrafish Immunohistochemistry

Whole mount IHC using rabbit anti-LCP1 (1:200, Genetex GTX124420) was performed, as previously described (19). Antibody staining using rabbit anti-5HT (1:200, Immunostar 20080) was performed as described before (72). Zebrafish were fixed in 4% PFA, dehydrated in 70% ethanol, and processed with a blocking solution (0.2% BSA, 0.5% Triton X-100, 1% DMSO, 5% horse serum, PBS). They were incubated overnight with primary antibody at 4°C, followed by overnight anti-rabbit Cy3 (1:250, Jackson) incubation. Finally, they were stained with DAPI for 3 minutes and stored in PBS at 4°C.

#### Zebrafish Whole-mount *in situ* hybridization

Whole mount *in situ* hybridization (WISH) with Digoxyigenin (DIG)-labeled was performed using standard protocols, as described previously (73). In short, 5 dpf wildtype and *ret* mutant zebrafish were fixed in 4% PFA, dehydrated in a graded methanol series (0%/25%/50%/100%), and subsequent rehydrated. To permeabilize the zebrafish, they underwent a 15 minute protein K treatment at 28,5°C, followed by refixation, and prehybridization at 65°C. Hybridization with denatured probes occurred overnight at 65°C, followed by post-hybridization washes. Blocking preceded an overnight incubation with Roche anti-DIG antibody. Subsequent washes, staining in 5μL NBT/BCIP per mL, and a stop solution were performed. After clearing in 50% glycerol/PBST, embryos were stored in glycerol.

*Fabp2* probe was designed with the following sequence: Antisense probe: aattaatacgactcactataGGCTTTAGGGCTGCCAATCATTAAAGC. Sense probe: TACGAGAAGTTCATGGAACAAATGG. Images were taken using the Olympus SZX1 microscope.

#### Zebrafish Whole-mount fluorescent *in situ* hybridization

Whole-mount fluorescent *in situ* hybridization (FISH) was performed as described before (24) The RNAscope Multiplex Fluorescence Reagent Kit v2 Assay (Advanced Cell Diagnostics, Bio-Techne) was used according to the manufacturers’ instructions. A custom-made probe for dr-*best4* C1 was used (NPR-0035110, Advanced Cell Diagnostics, Bio-Techne). Opal 570 dye (Akoya Biosciences) was used for channel development.

#### Live imaging zebrafish

Confocal imaging was performed using the SP5 intravital microscope with a 20x water-dipping lens, as previously described (74). Imaging of the 5 dpf tg(*phox2bb*:GFP; *kdrl*:mcherry) zebrafish was performed using the 488 nm and 561 nm laser. Briefly, zebrafish were anesthetized with 0.016% MS-222 in HEPES-buffered E3 and securely positioned in a row within low-melting agarose for stability during imaging. The imaging dish containing the embedded zebrafish, was filled with HEPES-buffered E3 containing 0.016% MS-222.

#### Quantification of IHC and FISH stainings in zebrafish

The number of immune cells, endothelial cells, ECCs and BEST4+ enterocytes were manually counted through Z-stack projections using FIJI (75). Length of the gut was measured using the straight-line tool. The number of counted cells was normalized to the gut length.

#### Human IHC and confocal microcopy

Immunofluorescence for CD45 (1:200 #AMAb90518, Atlas Antibodies) and 5-HT (1:200 #20080, Immunostar) was performed on pediatric tissue blocks obtained from the Department of Pathology of the Erasmus Medical Center, as previously described (25). Shortly, 4 µm sections were cut, deparaffinized using xylene and a graded ethanol series, followed by a pressure cooker treatment in 0.01M sodium citrate. Primary antibody was incubated overnight at 4°C and secondary antibody was incubated for 1 hour at room temperature. Anti-rabbit Cy3 (1:200, Jackson) was used for the 5-HT stainings and anti-mouse Alexa 488 (1:200, Jackson) for the CD45 stainings. Finally, DAPI staining was applied for 1 minute. The slides were then rinsed and mounted using a fluorescence mounting medium. Confocal imaging for 5-HT, CD45 and FABP2 was performed using the Stellaris confocal microscope with both 20x and 40x lenses. BEST4 stainings were performed by automated IHC using the Ventana Benchmark ULTRA (Ventana Medical Systems Inc.). Sequential 4 µm thick (FFPE) sections were stained using optiview (OV) (#760-700, Ventana). In brief, following deparaffinization and heat-induced antigen retrieval with CC1 (#950-500, Ventana) for 32 minutes, the tissue samples were incubated with BEST4 antibody (1:100 #HPA058564, Atlas Antibodies) for 32 minutes at 37°C. Incubation was followed by optiview detection and hematoxylin II counter stain for 8 minutes, followed by DAB staining, according to the manufactures instructions (Ventana). BEST4 stained slides were scanned using the Nanozoomer slide scanner (Hamamatsu) at the Pathology department of the Erasmus Medical Center.

#### Image analysis and quantification of human IHC stainings

Image analysis and quantification of human IHC stainings involved manual counting of 5-HT cells per crypt using FIJI (75). For CD45, a FIJI threshold was applied, and counts were determined per square, with 5 equal squares analyzed per slide. QuPath was used for BEST4 staining quantifications (76). Epithelial regions were selected to facilitate automated cell counting. Only the cone-shaped cells present in the epithelium were considered as BEST4+ cells.

#### Statistical analysis

For proportion testing of the scRNA-seq datasets of wildtype and *ret* mutant zebrafish, R scProportionTest package was used. Significant changes were determined with a permutation test to calculate the false discovery rate (FDR) and log_2_ fold change for each cluster. A confidence interval for the magnitude difference was returned via bootstrapping. FDR < 0.05 and absolute log_2_ fold change > 0.58 was considered significant.

Prior to comparing the mean cell counts between wildtype and *ret* mutant zebrafish, as well as for each staining comparing human controls and HSCR patients, an F-test was conducted to assess the equality of variances between the control and HSCR groups. If the F-test result indicated no significant difference in variances (p ≥ 0.05), Student’s t-test was used to compare group means. Conversely, if the F-test revealed a significant difference in variances (p < 0.05), Welch’s t-test was applied to account for unequal variances.

## Acknowledgments

We thank Iain Shepherd (Department of Biology, Emory University, Atlanta, USA) and Kaushal Parikh (Department of Gastroenterology and Hepatology, Erasmus University Medical Center, Rotterdam, NL) for their valuable feedback on the manuscript. We also thank Remko Hoogenboezem (Department of Hematology) for processing our raw single-cell data, and the optical imaging center (OIC) of the Erasmus University Medical Center (Rotterdam, NL) for assistance with confocal microscopy. This work was funded by the Friends of the Sophia Foundation (SSWO WAR-63).

## Author contributions

NJMK, LEK, RMWH, VM and MMA designed and planned the experiments; NJMK, LEK, JDW, JTMZ, MV, BMdG and DS prepared and executed the experiments; NJMK, WZ, LEK and EB supervised, prepared and analysed the single cell RNA sequencing data; TPPvdB provided human intestinal biopsies; DH, CEJS and RMWH provided clinical information; VM, EdP and MMA provided supervision and guidance; NJMK, WZ, LEK, VM and MMA interpreted the data and wrote the manuscript.

## Inclusion and diversity

We support inclusive, diverse, and equitable conduct of research.

## Declaration of generative AI and AI-assisted technologies

During the preparation of this work the author(s) used ChatGPT in order to improve language and grammar of the manuscript. After using this tool/service, the author(s) reviewed and edited the content as needed and take(s) full responsibility for the content of the publication.

## Supplemental information

Figure S1 and Table S1.

Data S1. Table showing the marker expression for all the clusters in the total intestinal dataset, related to Figure 1C.

Data S2. Table showing the marker expression for ENS subclusters, related to Figure 2B.

Data. S3. Table showing the marker expression for immune subclusters, related to Figure 3B.

Data S4. Table showing the marker expression for epithelial subclusters, related to Figure 4B.

Data S5. Table showing the marker expression for ECM subclusters, related to Figure 6B.

## Data and materials availability

- The raw and processed data used for this study will be available in NCBIs Gene Expression Omnibus through the GEO series accession number GSE225510 and GSE271622. The data can be accessed using the secure token afelugucjfgfhyb.
- This paper does not report original code.
- Any additional information required to reanalyze the data reported in this paper is available upon request.

## References

1. Wallace AS, Burns AJ. Development of the enteric nervous system, smooth muscle and interstitial cells of Cajal in the human gastrointestinal tract. Cell Tissue Res. 2005;319(3):367–82.

2. Nagy N, Goldstein AM. Enteric nervous system development: A crest cell’s journey from neural tube to colon. Semin Cell Dev Biol. 2017;66:94–106.

3. Goldstein AM, Thapar N, Karunaratne TB, De Giorgio R. Clinical aspects of neurointestinal disease: Pathophysiology, diagnosis, and treatment. Dev Biol. 2016;417(2):217–28.

4. Parisi MA, Kapur RP. Genetics of Hirschsprung disease. Curr Opin Pediatr. 2000;12(6):610–7.

5. Badner JA, Sieber WK, Garver KL, Chakravarti A. A genetic study of Hirschsprung disease. Am J Hum Genet. 1990;46(3):568–80.

6. Whitehouse FR, Kernohan JW. Myenteric plexus in congenital megacolon; study of 11 cases. Arch Intern Med (Chic). 1948;82(1):75–111.

7. Natarajan D, Marcos-Gutierrez C, Pachnis V, de Graaff E. Requirement of signalling by receptor tyrosine kinase RET for the directed migration of enteric nervous system progenitor cells during mammalian embryogenesis. Development. 2002;129(22):5151–60.

8. Taraviras S, Marcos-Gutierrez CV, Durbec P, Jani H, Grigoriou M, Sukumaran M, et al. Signalling by the RET receptor tyrosine kinase and its role in the development of the mammalian enteric nervous system. Development. 1999;126(12):2785–97.

9. Attie T, Pelet A, Edery P, Eng C, Mulligan LM, Amiel J, et al. Diversity of RET proto-oncogene mutations in familial and sporadic Hirschsprung disease. Hum Mol Genet. 1995;4(8):1381–6.

10. Angrist M, Bolk S, Halushka M, Lapchak PA, Chakravarti A. Germline mutations in glial cell line-derived neurotrophic factor (GDNF) and RET in a Hirschsprung disease patient. Nat Genet. 1996;14(3):341–4.

11. Swenson O, Neuhauser EB, Pickett LK. New concepts of the etiology, diagnosis and treatment of congenital megacolon (Hirschsprung’s disease). Pediatrics. 1949;4(2):201–9.

12. Soave F. Hirschsprung’s Disease: A New Surgical Technique. Arch Dis Child. 1964;39(204):116–24.

13. Duhamel B. [New operation for congenital megacolon: retrorectal and transanal lowering of the colon, and its possible application to the treatment of various other malformations] Une nouvelle operation pour le megacolon congenital: l’abaissement retro-rectal et trans-anal du colon et son application possible au traitement de quelques autres malformations. Presse Med (1893). 1956;64(95):2249-50.

14. M E, S A. Obstructive complications after pull-through for Hirschsprung’s disease: different causes and tailored management. Annals of Pediatric Surgery. 2019.

15. Hackam DJ, Filler RM, Pearl RH. Enterocolitis after the surgical treatment of Hirschsprung’s disease: risk factors and financial impact. J Pediatr Surg. 1998;33(6):830–3.

16. Moore SW, Albertyn R, Cywes S. Clinical outcome and long-term quality of life after surgical correction of Hirschsprung’s disease. J Pediatr Surg. 1996;31(11):1496–502.

17. Vincent E, Chatterjee S, Cannon GH, Auer D, Ross H, Chakravarti A, et al. Ret deficiency decreases neural crest progenitor proliferation and restricts fate potential during enteric nervous system development. Proc Natl Acad Sci U S A. 2023;120(34):e2211986120.

18. Wallace KN, Akhter S, Smith EM, Lorent K, Pack M. Intestinal growth and differentiation in zebrafish. Mech Dev. 2005;122(2):157–73.

19. Heanue TA, Boesmans W, Bell DM, Kawakami K, Vanden Berghe P, Pachnis V. A Novel Zebrafish ret Heterozygous Model of Hirschsprung Disease Identifies a Functional Role for mapk10 as a Modifier of Enteric Nervous System Phenotype Severity. PLoS Genet. 2016;12(11):e1006439.

20. Uribe RA. Genetic regulation of enteric nervous system development in zebrafish. Biochem Soc Trans. 2024;52(1):177–90.

21. Shepherd IT, Pietsch J, Elworthy S, Kelsh RN, Raible DW. Roles for GFRalpha1 receptors in zebrafish enteric nervous system development. Development. 2004;131(1):241–9.

22. Kuil LE, Chauhan RK, Cheng WW, Hofstra RMW, Alves MM. Zebrafish: A Model Organism for Studying Enteric Nervous System Development and Disease. Front Cell Dev Biol. 2020;8:629073.

23. Nechiporuk A, Linbo T, Poss KD, Raible DW. Specification of epibranchial placodes in zebrafish. Development. 2007;134(3):611–23.

24. Kuil LE, Kakiailatu NJM, Windster JD, Bindels E, Zink JTM, van der Zee G, et al. Unbiased characterization of the larval zebrafish enteric nervous system at a single cell transcriptomic level. iScience. 2023;26(7):107070.

25. Windster JD, Kuil LE, Kakiailatu NJM, Antanaviciute A, A. S, C. M, et al. Human enteric glia diversity in health and disease: new avenues for the treatment of Hirschsprung disease. BioRxiv. 2023.

26. Page DM, Wittamer V, Bertrand JY, Lewis KL, Pratt DN, Delgado N, et al. An evolutionarily conserved program of B-cell development and activation in zebrafish. Blood. 2013;122(8):e1–11.

27. Kaminski S, Hermann-Kleiter N, Meisel M, Thuille N, Cronin S, Hara H, et al. Coronin 1A is an essential regulator of the TGFbeta receptor/SMAD3 signaling pathway in Th17 CD4(+) T cells. J Autoimmun. 2011;37(3):198–208.

28. Guan J, Han S, Wu J, Zhang Y, Bai M, Abdullah SW, et al. Ribosomal Protein L13 Participates in Innate Immune Response Induced by Foot-and-Mouth Disease Virus. Front Immunol. 2021;12:616402.

29. Dai Y, Wu S, Cao C, Xue R, Luo X, Wen Z, et al. Csf1rb regulates definitive hematopoiesis in zebrafish. Development. 2022;149(16).

30. Macaulay IC, Svensson V, Labalette C, Ferreira L, Hamey F, Voet T, et al. Single-Cell RNA-Sequencing Reveals a Continuous Spectrum of Differentiation in Hematopoietic Cells. Cell Rep. 2016;14(4):966–77.

31. Hu C, Zhang N, Hong Y, Tie R, Fan D, Lin A, et al. Single-cell RNA sequencing unveils the hidden powers of zebrafish kidney for generating both hematopoiesis and adaptive antiviral immunity. Elife. 2024;13.

32. Zhou Q, Zhao C, Yang Z, Qu R, Li Y, Fan Y, et al. Cross-organ single-cell transcriptome profiling reveals macrophage and dendritic cell heterogeneity in zebrafish. Cell Rep. 2023;42(7):112793.

33. Hernandez PP, Strzelecka PM, Athanasiadis EI, Hall D, Robalo AF, Collins CM, et al. Single-cell transcriptional analysis reveals ILC-like cells in zebrafish. Sci Immunol. 2018;3(29).

34. Willms RJ, Jones LO, Hocking JC, Foley E. A cell atlas of microbe-responsive processes in the zebrafish intestine. Cell Reports. 2022;38(5):110311.

35. Lechuga S, Marino-Melendez A, Davis A, Zalavadia A, Khan A, Longworth MS, et al. Coactosin-like protein 1 regulates integrity and repair of model intestinal epithelial barriers via actin binding dependent and independent mechanisms. Front Cell Dev Biol. 2024;12:1405454.

36. Li Y, Li X, Geng C, Guo Y, Wang C. Somatostatin receptor 5 is critical for protecting intestinal barrier function in vivo and in vitro. Mol Cell Endocrinol. 2021;535:111390.

37. Le H, Lie KK, Giroud-Argoud J, Ronnestad I, Saele O. Effects of Cholecystokinin (CCK) on Gut Motility in the Stomachless Fish Ballan Wrasse (Labrus bergylta). Front Neurosci. 2019;13:553.

38. Li HJ, Ray SK, Pan N, Haigh J, Fritzsch B, Leiter AB. Intestinal Neurod1 expression impairs paneth cell differentiation and promotes enteroendocrine lineage specification. Sci Rep. 2019;9(1):19489.

39. Hill ME, Asa SL, Drucker DJ. Essential requirement for Pax6 in control of enteroendocrine proglucagon gene transcription. Mol Endocrinol. 1999;13(9):1474–86.

40. Rehfeld JF. Cholecystokinin-From Local Gut Hormone to Ubiquitous Messenger. Front Endocrinol (Lausanne). 2017;8:47.

41. Terry NA, Walp ER, Lee RA, Kaestner KH, May CL. Impaired enteroendocrine development in intestinal-specific Islet1 mouse mutants causes impaired glucose homeostasis. Am J Physiol Gastrointest Liver Physiol. 2014;307(10):G979–91.

42. Stijnen P, Ramos-Molina B, O’Rahilly S, Creemers JW. PCSK1 Mutations and Human Endocrinopathies: From Obesity to Gastrointestinal Disorders. Endocr Rev. 2016;37(4):347–71.

43. Yu Y, Yang W, Li Y, Cong Y. Enteroendocrine Cells: Sensing Gut Microbiota and Regulating Inflammatory Bowel Diseases. Inflamm Bowel Dis. 2020;26(1):11–20.

44. Wu Q, Li W, You C. The regulatory roles and mechanisms of the transcription factor FOXF2 in human diseases. PeerJ. 2021;9:e10845.

45. Quest AF, Gutierrez-Pajares JL, Torres VA. Caveolin-1: an ambiguous partner in cell signalling and cancer. J Cell Mol Med. 2008;12(4):1130–50.

46. Aimes RT, Quigley JP. Matrix metalloproteinase-2 is an interstitial collagenase. Inhibitor-free enzyme catalyzes the cleavage of collagen fibrils and soluble native type I collagen generating the specific 3/4- and 1/4-length fragments. J Biol Chem. 1995;270(11):5872-6.

47. Zhang Y, Hu W. NFkappaB signaling regulates embryonic and adult neurogenesis. Front Biol (Beijing). 2012;7(4).

48. Meffert MK, Chang JM, Wiltgen BJ, Fanselow MS, Baltimore D. NF-kappa B functions in synaptic signaling and behavior. Nat Neurosci. 2003;6(10):1072–8.

49. Simmons LJ, Surles-Zeigler MC, Li Y, Ford GD, Newman GD, Ford BD. Regulation of inflammatory responses by neuregulin-1 in brain ischemia and microglial cells in vitro involves the NF-kappa B pathway. J Neuroinflammation. 2016;13(1):237.

50. Liu T, Zhang L, Joo D, Sun SC. NF-kappaB signaling in inflammation. Signal Transduct Target Ther. 2017;2:17023-.

51. Elkrewi EZ, Al Abdulqader AA, Khasanov R, Maas-Omlor S, Boettcher M, Wessel LM, et al. Role of Inflammation and the NF-kappaB Signaling Pathway in Hirschsprung’s Disease. Biomolecules. 2024;14(8).

52. Dariel A, Grynberg L, Auger M, Lefevre C, Durand T, Aubert P, et al. Analysis of enteric nervous system and intestinal epithelial barrier to predict complications in Hirschsprung’s disease. Sci Rep. 2020;10(1):21725.

53. Perea D, Guiu J, Hudry B, Konstantinidou C, Milona A, Hadjieconomou D, et al. Ret receptor tyrosine kinase sustains proliferation and tissue maturation in intestinal epithelia. EMBO J. 2017;36(20):3029–45.

54. Shepherd A, Feinstein L, Sabel S, Rastelli D, Mezhibovsky E, Matthews L, et al. RET Signaling Persists in the Adult Intestine and Stimulates Motility by Limiting PYY Release from Enteroendocrine Cells. bioRxiv. 2022.

55. Onizawa M, Nagaishi T, Kanai T, Nagano K, Oshima S, Nemoto Y, et al. Signaling pathway via TNF-alpha/NF-kappaB in intestinal epithelial cells may be directly involved in colitis-associated carcinogenesis. Am J Physiol Gastrointest Liver Physiol. 2009;296(4):G850–9.

56. Jones LG, Vaida A, Thompson LM, Ikuomola FI, Caamano JH, Burkitt MD, et al. NF-kappaB2 signalling in enteroids modulates enterocyte responses to secreted factors from bone marrow-derived dendritic cells. Cell Death Dis. 2019;10(12):896.

57. Henderson J, O’Reilly S. The emerging role of metabolism in fibrosis. Trends Endocrinol Metab. 2021;32(8):639–53.

58. Romero Y, Aquino-Galvez A. Hypoxia in Cancer and Fibrosis: Part of the Problem and Part of the Solution. Int J Mol Sci. 2021;22(15).

59. Sakai T, Choo YY, Sato O, Ikebe R, Jeffers A, Idell S, et al. Myo5b Transports Fibronectin-Containing Vesicles and Facilitates FN1 Secretion from Human Pleural Mesothelial Cells. Int J Mol Sci. 2022;23(9).

60. Dou F, Liu Q, Lv S, Xu Q, Wang X, Liu S, et al. FN1 and TGFBI are key biomarkers of macrophage immune injury in diabetic kidney disease. Medicine (Baltimore). 2023;102(45):e35794.

61. He S, Wang J, Huang Y, Kong F, Yang R, Zhan Y, et al. Intestinal fibrosis in aganglionic segment of Hirschsprung’s disease revealed by single-cell RNA sequencing. Clinical and Translational Medicine. 2023;13(2).

62. Bertrand JY, Chi NC, Santoso B, Teng S, Stainier DY, Traver D. Haematopoietic stem cells derive directly from aortic endothelium during development. Nature. 2010;464(7285):108-11.

63. Kuil LE, Kakiailatu NJM, Windster JD, Bindels E, Zink JTM, van der Zee G, et al. Unbiased characterization of the larval zebrafish enteric nervous system at a single cell transcriptomic level. iScience. 2023;26(7).

64. Kakiailatu NJM, Kuil LE, Bindels E, Zink JTM, Vermeulen M, Melotte V, et al. Gut Isolation from Zebrafish Larvae for Single-cell RNA Sequencing. J Vis Exp. 2023(201).

65. Hao Y, Stuart T, Kowalski MH, Choudhary S, Hoffman P, Hartman A, et al. Dictionary learning for integrative, multimodal and scalable single-cell analysis. Nat Biotechnol. 2024;42(2):293–304.

66. Stuart T, Butler A, Hoffman P, Hafemeister C, Papalexi E, Mauck WM, 3rd, et al. Comprehensive Integration of Single-Cell Data. Cell. 2019;177(7):1888–902 e21.

67. Miller SA, Policastro RA, Sriramkumar S, Lai T, Huntington TD, Ladaika CA, et al. LSD1 and Aberrant DNA Methylation Mediate Persistence of Enteroendocrine Progenitors That Support BRAF-Mutant Colorectal Cancer. Cancer Res. 2021;81(14):3791–805.

68. Dolgalev I. msigdbr: MSigDB Gene Sets for Multiple Organisms in a Tidy Data Format. 7.5.1.9001 ed: R package; 2024.

69. Korotkevich G, Sukhov V, Budin N, Shpak B, Artyomov MN, Sergushichev A. 2021.

70. Trapnell C, Cacchiarelli D, Grimsby J, Pokharel P, Li S, Morse M, et al. The dynamics and regulators of cell fate decisions are revealed by pseudotemporal ordering of single cells. Nat Biotechnol. 2014;32(4):381–6.

71. Qiu X, Hill A, Packer J, Lin D, Ma YA, Trapnell C. Single-cell mRNA quantification and differential analysis with Census. Nat Methods. 2017;14(3):309–15.

72. Hyland C, Mfarej M, Hiotis G, Lancaster S, Novak N, Iovine MK, et al. Impaired Cx43 gap junction endocytosis causes morphological and functional defects in zebrafish. Mol Biol Cell. 2021;32(20):ar13.

73. Thisse C, Thisse B. High-resolution in situ hybridization to whole-mount zebrafish embryos. Nat Protoc. 2008;3(1):59–69.

74. Kuil LE, Oosterhof N, Ferrero G, Mikulášová T, Hason M, Dekker J, et al. Zebrafish macrophage developmental arrest underlies depletion of microglia and reveals Csf1r-independent metaphocytes. eLife. 2020;9:e53403.

75. Schneider CA, Rasband WS, Eliceiri KW. NIH Image to ImageJ: 25 years of image analysis. Nat Methods. 2012;9(7):671–5.

76. Bankhead P, Loughrey MB, Fernandez JA, Dombrowski Y, McArt DG, Dunne PD, et al. QuPath: Open source software for digital pathology image analysis. Sci Rep. 2017;7(1):16878.

77. Liberzon A, Birger C, Thorvaldsdottir H, Ghandi M, Mesirov JP, Tamayo P. The Molecular Signatures Database (MSigDB) hallmark gene set collection. Cell Syst. 2015;1(6):417–25.

