## Supplemental Figure 1 for "Impact of Enteric Neuronal Loss on Intestinal Cell Composition"

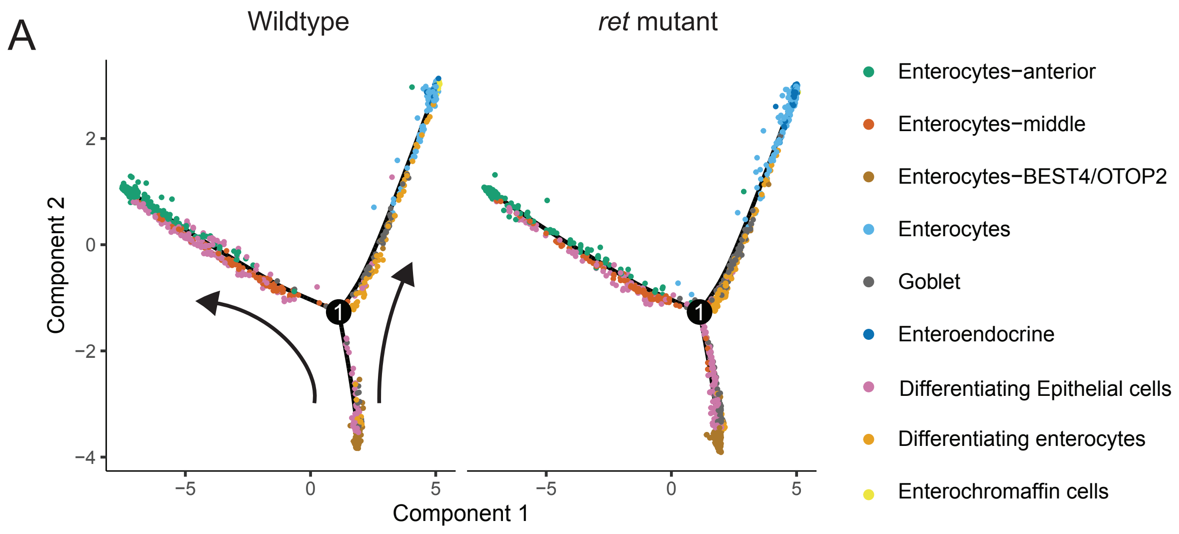


**Supplementary Figure 1.**

Pseudotime trajectories showing the developmental track of epithelial cells in wildtype (left panel) and *ret* mutant (right panel) zebrafish intestines. Each dot represents an individual epithelial cell, color-coded by cell type corresponding to Figure 4A. The black arrows indicate the inferred developmental trajectory.
