## Supplemental Table 1 for "Impact of Enteric Neuronal Loss on Intestinal Cell Composition"

| Sample ID | Sex | Age at surgery | Indication for surgery | Bowel region | *RET* Mutation |
| --- | --- | --- | --- | --- | --- |
| CTRL1 | Female | 5 months | Rectum prolaps | Rectosigmoid | - |
| CTRL2 | Male | 15 days | Milk Curd | Rectum | - |
| CTRL3 | Female | 5y + 3 months | ARM | Rectosigmoid | - |
| CTRL4 | Male | 3 months | NEC | Colon ascendens | - |
| CTRL5 | Male | 3 months | NEC | Colon ascendens | - |
| HSCR1 | Male | 10 months | HSCR | Rectosigmoid | c.988C>T, p.(Arg330Trp) |
| HSCR2 | Male | 7.5 months | HSCR | Rectosigmoid | c.229C>T, p.(Arg77Cys) |
| HSCR3 | Male | 6 months | HSCR | Rectosigmoid | c.868-7C>G |
| HSCR4 | Male | 7 months | HSCR | Rectosigmoid | c.692G>A, p.(Arg231His) |
| HSCR5 | Male | 3 months | HSCR | Rectosigmoid | c.2690G>A, p.Arg897Gln |

**Supplementary Table 1 – Patient diagnosis and major characteristics**
